# Evolution is All You Need in Promoter Design and Optimization

**DOI:** 10.1101/2023.11.18.567645

**Authors:** Ruohan Ren, Hongyu Yu, Jiahao Teng, Sihui Mao, Zixuan Bian, Yangtianze Tao, Stephen S.-T. Yau

**Affiliations:** Zhili College, Tsinghua University, Beijing, China; Department of Mathematical Sciences, Tsinghua University, Beijing, China; School of Life Sciences, Tsinghua University, Beijing, China; Weiyang College, Tsinghua University, Beijing, China; Yanqi Lake Beijing Institute of Mathematical Science and Applications, Beijing, China

## Abstract

Predicting the strength of promoters and guiding their directed evolution is a crucial task in synthetic biology. This approach significantly reduces the experimental costs in conventional promoter engineering. Previous studies employing machine learning or deep learning methods have shown some success in this task, but their outcomes were not satisfactory enough, primarily due to the neglect of evolutionary information. In this paper, we introduce the Chaos-Attention net for Promoter Evolution (CAPE) to address the limitations of existing methods. We comprehensively extract evolutionary information within promoters using chaos game representation and process the overall information with DenseNet and Transformer. Our model achieves state-of-the-art results on two kinds of distinct tasks. The incorporation of evolutionary information enhances the model’s accuracy, with transfer learning further extending its adaptability. Furthermore, experimental results confirm CAPE’s efficacy in simulating in silico directed evolution of promoters, marking a significant advancement in predictive modeling for prokaryotic promoter strength. Our paper also presents a user-friendly website for the practical implementation of in silico directed evolution on promoters.

## 1 Introduction

In the field of synthetic biology, precise characterization of regulatory elements is of paramount importance for the design of synthetic gene circuits [1] [2]. Such characterization can significantly advance various critical domains, including pharmaceutical synthesis [3] [4], metabolic engineering [5], and material production [6] [7]. Among the various regulatory elements, promoters play a pivotal role in synthetic biology [8], as they exert significant control over the expression level of downstream genes [9] [10]. Therefore, identifying a promoter with the appropriate strength is crucial for constructing expression vectors, and the optimization of promoter sequences is a key task in synthetic biology.

Conventional promoter engineering relies on experimental techniques for identifying suitable promoters, including mutagenesis [11] [12], sequence combinations [13], etc. One commonly adopted method involves random mutagenesis of promoters through error-prone PCR, followed by the selection of mutants with increased strength [14]. This iterative process is often referred to as the directed evolution of promoters. Nevertheless, experimental methods are frequently characterized by high levels of unpredictability, labor intensiveness, and inherent stochasticity in the results.

The development of artificial intelligence has created the foundation for in silico directed evolution of promoters, with a key prerequisite being the establishment of an accurate regression model that correlates promoter sequences with their strengths. There have been some related research on computational models for prokaryotic promoters, including machine learning/deep learning models. However, many models are used to identify whether a given sequence can serve as a promoter [15] [16] [17]. For promoter strength prediction, most of the existing models are classification models, used to predict whether a promoter is strong or weak [18] [19] [20]. Currently, there is still a lack of accurate regression models in this regard. It is worth mentioning that Wang et al. combined a deep generative model with a predictive model to preselect the most promising synthetic promoters [21]. However, due to the difficulty of avoiding noise interference, the Pearson correlation coefficient (PCC) of their CNN-based prediction model was around 0.25, suggesting a pressing need for substantial improvement.

Understanding the evolutionary history of corresponding promoters plays a crucial role when aiming for directed promoter evolution. The significance of coevolutionary information in deep learning models has been successfully demonstrated in fields such as protein structure prediction [22] and proteinprotein interaction prediction [23]. However, in the field of promoter design, suitable coevolutionary information features have yet to be successfully applied. The limited availability of promoter data poses a challenge for feature extraction using traditional alignment algorithms, as finding an adequate number of suitably similar promoters proves to be difficult. Leveraging the alignment-free chaos game representation [24] allows us to extract the inherent coevolutionary information within promoters with only local or moderate similarity, offering valuable support to enhance the model’s effectiveness.

In this paper, we propose the Chaos-Attention net for Promoter Evolution (CAPE), which features the incorporation of chaos game representation and the utilization of DenseNet [25] and Transformer [26]. CAPE is a highly accurate regression model that establishes correlations between promoter sequences and their strengths, leading to state-of-the-art (SOTA) performance. Through the coevolutionary information extracted from promoter sequences, our deep learning model achieved a PCC of 0.56 on the dataset from [27], significantly surpassing 0.25 obtained by Wang et al. [21] and 0.29 obtained by the predictor of DeepSEED [28] on the same dataset. Furthermore, we implemented transfer learning to enhance the model’s adaptability to a range of downstream tasks. For example, when applied to predict the strength of trc promoters, our model achieved an R-squared (R2) value of 0.77, signifying a substantial improvement over Zhao et al.’s method (0.53) [29] and six EVMP-based algorithms (0.65 for the best one) [30]. This underscores the considerable superiority of our model structure. Finally, we conducted directed evolution experiments on a series of promoters, and the results indicate that our model is indeed capable of efficiently evolving promoters.

In summary, we harnessed coevolutionary information to construct a deep learning model CAPE, which enabled us to attain SOTA performance in predicting prokaryotic promoter strength. We confirmed the model’s effectiveness and wide applicability in simulating the directed evolution of promoters in silico through biological experiments. We also developed a website for convenient implementation of directed evolution for promoters.

## 2 Results

### 2.1 Overview of the Model Architecture

The architecture of CAPE is as follows (Figure 1).

**Figure 1:**
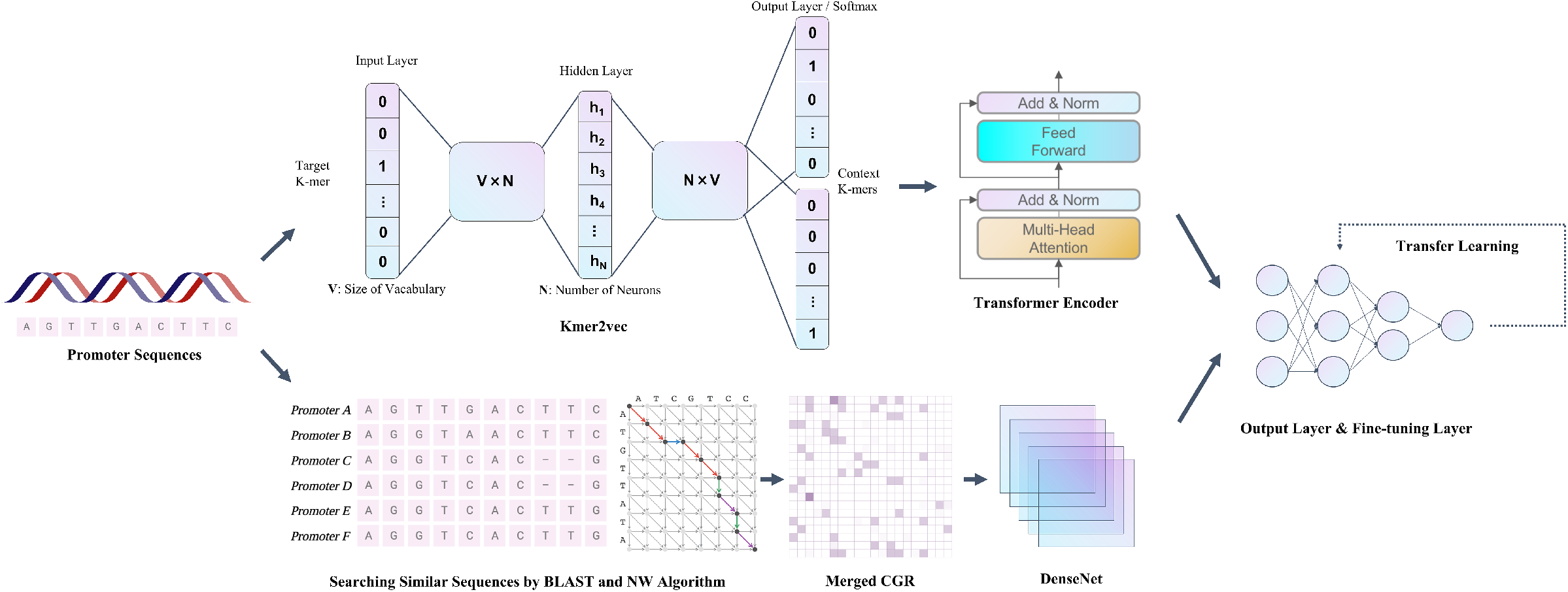
Overview of the Model Architecture

First, we employed BLAST [31] and the Needleman-Wunsch (NW) algorithm [32] to search for sequences exhibiting a certain level of similarity with the target promoter within a prokaryotic promoter database [33]. Subsequently, we applied a novel method firstly introduced in this paper, referred to as merged CGR, to convert the promoter sequence into image data capturing evolutionary information. Alongside image information, we applied the kmer2vec method [34] to extract textual information from the promoter sequences.

The above two types of information will be input into two different deep learning networks, namely DenseNet [25] and Transformer [26], respectively. The results processed by both models are fed into a fully connected network for integration. Finally, our model can output the predicted strengths of given promoter sequences. Moreover, we introduced a fine-tuning network for transfer learning, which enhances the model’s ability to adapt to various downstream tasks.

### 2.2 Modules of the Model

Here, we briefly introduce the structures of Merged CGR, DenseNet, Transformer, and the fully connected networks. For further details about the modules, please refer to Section 4.

Traditional Chaos Game Representation (CGR) method [24] transforms the sequence into a series of points within a unit square. Merged CGR is an extension of this method. Initially, the point series is further pixelated to create a matrix for ease of summation. Subsequently, matrices derived from the sequence and its related sequences are weighted and summed based on their similarity, resulting in the Merged CGR matrix. This process facilitates the integration of evolutionary and sequence information in the form of image data (Figure 2(a)). DenseNet is an extension of convolutional neural networks, enhancing the flow of information by directly concatenating layers to prevent information distortion. Due to limitations in the data size, we did not employ the deep network structure as in the original work. Instead, we embedded a relatively shallow network into the DenseNet architecture, as illustrated in Figure 2(b). In the figure, squares represent data, the “H” symbol signifies mappings within the network, and multiple lines converging towards the same “H” indicate that the input to this “H” is the concatenation result of these data.

**Figure 2:**
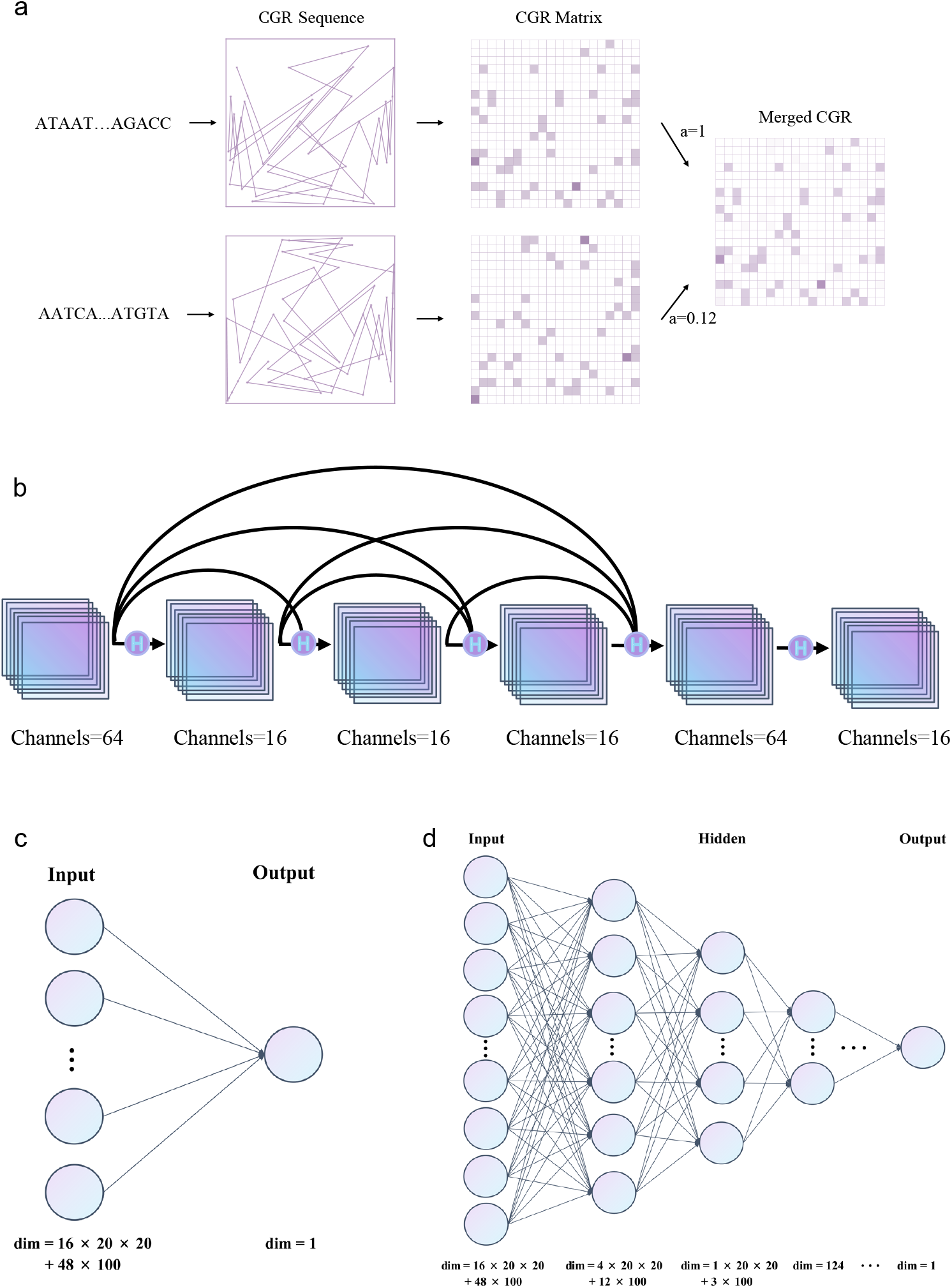
Modules of the Model. (a) Merged CGR. (b) DenseNet. (c) The original fully connected network. (d) The fully connected network for finetuning.

Transformer is a deep learning algorithm based on the multi-head attention mechanism designed for processing sequential information (Figure 1). Unlike other sequence-to-sequence tasks, here we are solely focused on regression. Therefore, we do not adopt the encoder-decoder structure; instead, we only utilize the encoder in traditional Transformer.

The fully connected network is utilized to combine the graphical evolutionary information processed by DenseNet and the sequential linguistic information processed by Transformer. A single-layer network suffices for the original model (Figure 2(c)). Throughout the fine-tuning process, we maintain the parameters of the preceding model as existing knowledge and solely replace the fully connected network for retraining. Therefore, to enhance flexibility, multiple layers of fully connected networks are employed (Figure 2(d)).

### 2.3 Performance on Two Prediction Tasks

To validate the effectiveness of CAPE, we conducted tests for two prediction tasks. The dataset for Task1, named dataset_Ecoli, comprises 11,884 promoter sequences along with their corresponding strengths. (Details are shown in section 4.) Each promoter is artificially defined as the 50 bp sequence preceding the transcription start site. These promoter sequences exhibit significant diversity, corresponding to different genes, allowing the use of dataset_Ecoli for the model’s general training. We randomly split the set in a 3:1 ratio, with 3/4 as the training set and 1/4 as the test set. In Figure 3(c), it can be observed that our split is random enough, resulting in a reasonably consistent distribution of promoter strengths between the training and test sets. Our model achieved a PCC of 0.56 on the test set of Task1, surpassing the 0.25 reported by Wang et al. [21] and the 0.29 produced by the predictor of DeepSEED [28]. Our model’s performance is almost 2 times higher than that of the previous best-performing model (Figure 3(e)). The model’s substantial enhancement is impressive, significantly improving the capacity to extract information and avoid extensive noise. The scatter plots of the model’s predictions on the training and test sets are also depicted (Figure 3(a)).

**Figure 3:**
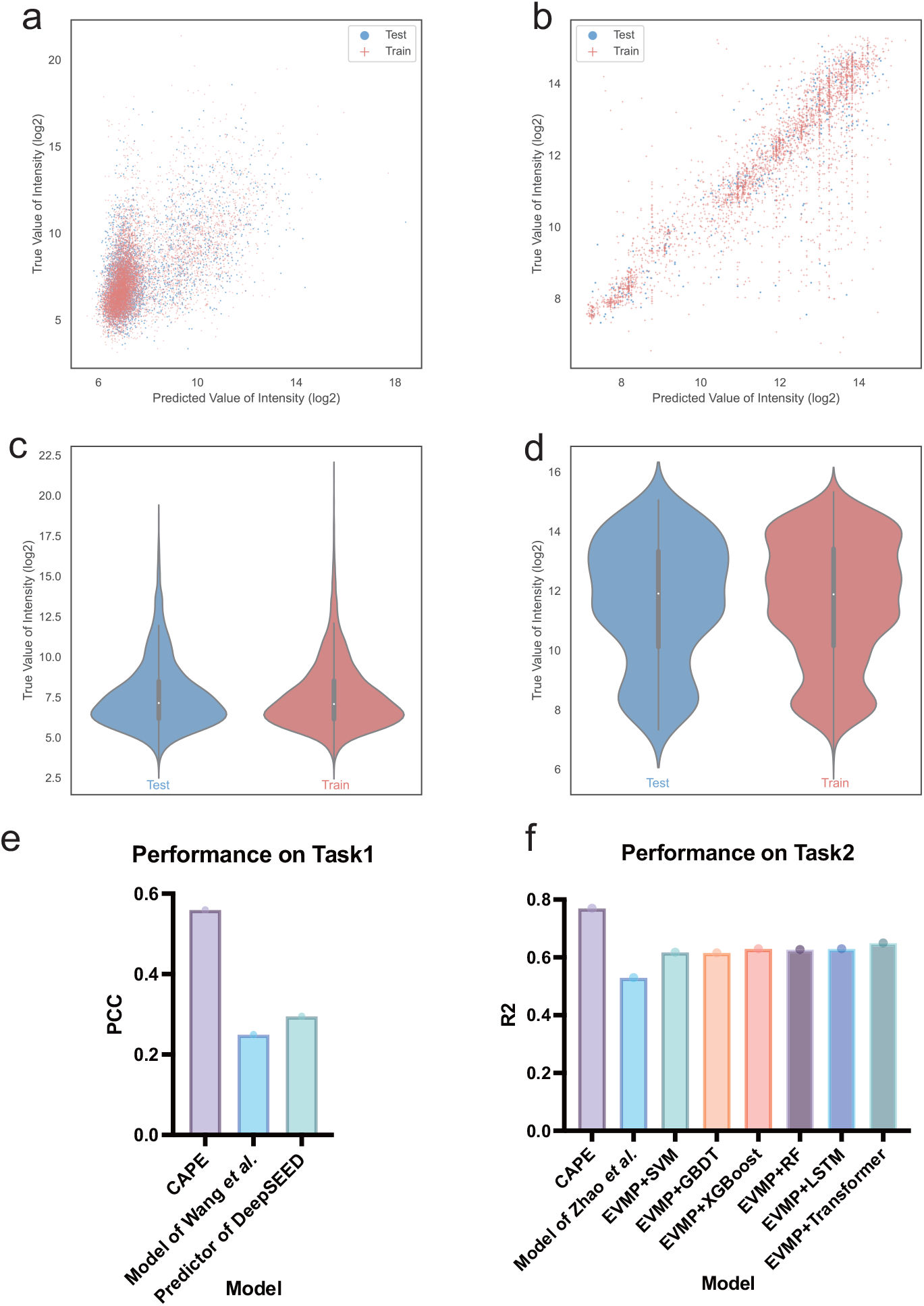
Performance of the Model. (a) The scatter plot for model prediction in Task1. (b) The scatter plot for model prediction in Task2. (c) The violin plot for true promoter strengths in Task1. (d) The violin plot for true promoter strengths in Task2. (e) PCC for the test set in Task1. (f) R2 for the test set in Task2.

In synthetic biology, Escherichia coli stands as an indisputable model organism in prokaryotes, and many other prokaryotic promoters demonstrate functionality in E. coli. Therefore, the model trained on dataset_Ecoli can aid in predicting prokaryotic promoter strength. Moreover, it can simulate mutational screening and directed evolution based on the predicted strengths.

As the strengths of promoters in dataset_Ecoli were measured by dRNAseq, this measurement method might introduce some noise. Currently, most promoter strength tests in experiments use fluorescent proteins (such as green fluorescent protein or monomeric Cherry fluorescent protein) as downstream reporters. We sought to understand whether our model is suitable for datasets using fluorescent protein strength as the promoter strength. Hence, we conducted Task2 test. We selected dataset_trc, containing 3,665 promoter sequences along with their corresponding strengths. (Details are shown in Section 4.) The trc promoter is a synthetic composite of trp and lac promoters [35] [36]. Though it can be induced, Zhao et al. [29] considered the trc promoter as a constitutive promoter for strength screening (by removing lacI). These 3,665 promoters are variant strains of the original trc promoter, displaying different strengths. Notably, Task2 differs significantly from Task1. The promoter sequences in Task1 dataset show considerable diversity, while those in Task2 have minor differences, demanding higher precision from the model in recognizing these distinctions. In addition, dataset_trc utilizes fluorescent protein strength to represent promoter strength, likely resulting in less noise.

To assess the model’s transfer learning capability, as the promoter length in Task1 was 50 bp, we needed to maintain consistency between both tasks. Here, we characterized the last 50 bp of the 74 bp trc promoter sequences in the dataset, as bases closer to the transcription start site typically have a more substantial influence. Similar to Zhao et al. [29], we randomly split the dataset in a 9:1 ratio, using 9/10 as the training set and 1/10 as the test set. Our random split did not affect the overall distribution of promoters with different strengths (Figure 3(d)). During training, we first transferred the model obtained from Task1, fixing all parameters before the fully connected network, as we believed these parameters contained a significant amount of information regarding prokaryotic promoters. Subsequently, we employed a pyramid-shaped fully connected layer as the fine-tuning network, replacing the original fully connected network. Only the parameters of the fine-tuning layer were modified during training.

The model achieved an R2 score of 0.77 on the test set. For fair comparison, we re-evaluated Zhao et al.’s method (based on XGBoost) [29] using their provided code and our identical dataset split. Their method scored an R2 of 0.53. We also evaluated six EVMP-based algorithms developed in [30] and obtained R2 values ranging from 0.62 to 0.65. This demonstrates that our model achieved an overall improvement of 18% compared to the previously best-performing model, showcasing exceptional transfer learning capability (Figure 3(f)). The scatter plot of the model’s predictions on the training and test sets is displayed (Figure 3(b)). Task2 validated the model’s transfer learning ability, showcasing that our model could utilize general information from Task1 to make more specific predictions such as Task2.

### 2.4 Experimental Results

To further validate the accuracy of CAPE, we proceeded to conduct verification experiments in the E. coli Stbl3 strain, utilizing constitutive promoter PnisA, as well as the heat-inducible promoter PL. Please note that the PnisA promoter is an nisin-induced promoter in Lactococcus lactis. However, due to its baseline expression after introduction into Escherichia coli, it is considered as a constitutive promoter in our research. The flowchart of the experimental process is shown in Figure 4(a). We used the original sequences as input to generate top-performing sequences through in silico directed evolution for each promoter. These sequences are detailed in Supplementary Table 2. Subsequently, we successfully employed molecular cloning techniques to construct the mutated promoters alongside the gene of mCherry fluorescent protein into pET28A and pBV220 plasmid vectors. The experimental results indicate that, fluorescence intensity analysis demonstrated a significant increase in promoter strength for the PnisA promoter. The highest fluorescence/OD600 (defined as unit fluorescence intensity), in comparison to the original sequence, increased to 234% (Figure 4(b)). Furthermore, 37.5% of the mutated promoters demonstrated enhanced expression strength when compared to the original promoter (Figure 4(b)). We also get similar results by single-cell fluorescence intensity detection in flow cytometry, and half of the mutated promoters (4/8) showed mCherry expression levels exceeding those of the original promoter, with the highest MFI (Median fluorescence intensity) surpassing the original promoter to 443% (Figure 4(b)). In addition to the constitutive promoters, we also obtained satisfactory results for the heat-inducible PL promoter. The induced fluorescence per OD600, compared to the wild-type, showed a maximum 497% increase, with 35.7% of the mutated promoters surpassing the wild-type expression level (Figure 4(c)). During flow cytometry detection, 57.1% of the mutant PL promoters showed higher mCherry expression than the original promoters, with the highest MFI exceeding the original promoter to 857% (Figure 4(c)).

**Figure 4:**
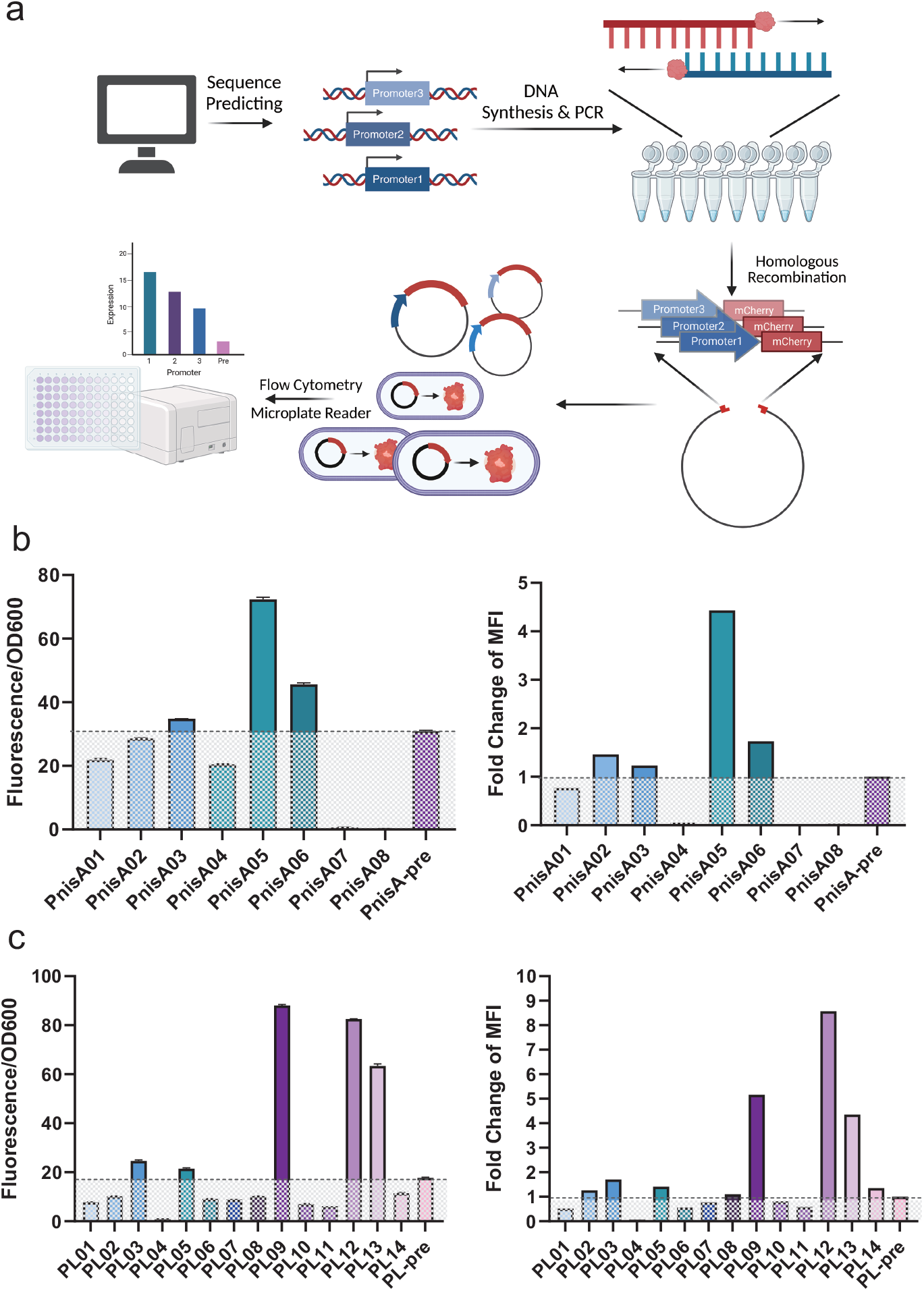
Experimental Results. (a) Flowchart of Experimental Process. (b) Results of PnisA Promoter. (c) Results of PL Promoter.

In conclusion, our biological experiments have preliminarily confirmed the effectiveness and reliability of our model, which can significantly enhance the expression strength of prokaryotic promoters.

## 3 Discussions

In summary, we have successfully built a powerful and useful tool, CAPE, to predict the strength of prokaryotic promoters, achieving SOTA performance in two distinct tasks. We have also validated the effectiveness and robustness of this tool through successful fluorescence expression assays. This deep learning model comprehensively understands the essence of promoters by leveraging coevolutionary information, holding significant importance. By grasping the biological essence of “evolution”, we have advanced breakthroughs in promoter design and optimization. Additionally, we have provided a website for users to freely utilize our tool.

From our perspective, CAPE can potentially play a pivotal role in several areas, including but not limited to the following:

1. In silico directed evolution and library construction of promoters: Our model accurately assesses promoter strength and can precisely identify minor differences in promoter sequences. Hence, we can conduct in silico directed evolution of promoters to obtain the desired sequences, facilitating our model’s crucial role in promoter optimization. Our model’s proficiency in directed evolution has been confirmed through multiple experiments. Additionally, should datasets containing mutated promoter strengths exist, integrating them with our model can refine and tailor more precise models, facilitating the development of experimental promoter libraries.
2. Enhancing the effectiveness of promoter generation models: Some research focuses on promoter generation models to produce synthetic promoters with improved functionality [21] [28] [37]. These models often face a selection step requiring precise prediction of promoter strength, where our model is likely to enhance the effectiveness of existing promoter generation models.
3. Reducing the cost of drug and metabolite production: Many drugs and metabolites are produced using model organisms like Escherichia coli [38] [39]. If our model is employed to direct the evolution of promoters, enhancing their strength, it could potentially reduce the manufacturing costs of related drugs and metabolites in the future.
4. Expansion from prokaryotic to eukaryotic organisms: Eukaryotic promoters are more complex compared to prokaryotic promoters and possess many other regulatory elements like enhancers [40] and silencers [41]. Currently, there’s no universally suitable precise model for eukaryotic organisms. However, in the future, we can primarily focus on the evolution of core eukaryotic promoters. With sufficiently large datasets suitable for deep learning training, our model structure could be further attempted in predicting the strength of eukaryotic core promoters.
5. Supporting epidemic control from a promoter standpoint: Recent public health crises such as SARS-CoV-2 have significantly impacted humanity. Various methods are attempting to help address this issue by focusing on diverse genomic elements, such as enhancer analysis [42] and codon adaptation analysis [43]. We can focus on this issue from a promoter standpoint. Our model’s structure may aid in advancing the evolution of eukaryotic promoters in the future, potentially reducing the cost of vaccines. In addition, we may also enhance the effectiveness of some SARS-CoV-2 detection methods involving promoters [44].
6. Other promoter strength-related issues: Numerous issues are related to prokaryotic and eukaryotic promoters. In agriculture, optimizing plant promoters can lead to increased gene expression, enhancing crop resistance, yield, and stress tolerance [45]. In environmental protection, regulating the gene expression of engineered bacteria aids in the metabolism and degradation of pollutants, contributing to environmental remediation and conservation efforts [46]. Studying promoters and gene expression regulation helps understand disease mechanisms, as abnormal promoter strength or regulatory elements might contribute to certain disease occurrences [47].

Of course, CAPE also has some aspects needing future improvement:

1. Currently, our model only accepts 50bp sequences for prediction. For larger promoters, using sequences closer to the transcription start site as a substitute has been proven effective. The sequence length limitation primarily stems from the dataset constraints. If more diverse datasets are integrated in the future, our model can further improve by overcoming this limitation through NLP methods such as padding [26].
2. Task1 dataset contains substantial noise, partly due to the use of dRNAseq for measurement and artificial definitions of promoters, resulting in the original model [21] reaching only around 0.25 PCC in performance. While we have significantly improved this to 0.56, the PCC is still not high enough. In the future, effectively integrating more DNA feature extraction methods might enhance our model’s performance. For instance, the K-mer Natural Vector method [48] has shown advanced effectiveness in phylogenetic analysis [49]. Additionally, methods like Z-curve [50], successful in promoter recognition [16], could also be considered.
3. Fine-tuning for Task2 depends on specific promoter strength data after mutation, requiring a substantial data accumulation. While our model’s predictive effectiveness achieves SOTA, it still requires experimental support. In the future, combining our model with active learning methods [51] may further save time and effort for experimentalists.
4. Despite conducting several experiments to validate the model’s effectiveness, in the future, we might consider adding more experimental data for further in-depth research.
5. Our method primarily focuses on predicting the strength of constitutive promoters and inducible promoters after induction. However, we observed some leakage in the mutated promoters before induction (see Supplementary Figure 1). This phenomenon can be attributed to sequence alterations that reduced the binding affinity of the original repressor protein with the promoter region, as anticipated. Nevertheless, we also identified mutated promoters that exhibited less pronounced leakage but a significant increase in expression levels after induction. This suggests that we may need to consider the specific kinetic dynamics of the inducible promoters in our subsequent work to enhance the model’s reliability. Designing a separate dual-task model based on our model structure could likely help address this issue. Furthermore, excessive leakage can potentially allow us to employ in silico directed evolution to transform inducible promoters into constitutive promoters, thereby achieving long-term stable gene expression.

## 4 Methods

### 4.1 Dataset

In our research, we utilized three datasets, which are as follows.

The first dataset, named dataset_pro, is derived from the PPD database and comprises 129,148 experimentally validated promoter sequences across 63 prokaryotic species [33]. We conducted sequence alignment within dataset_pro to identify similar promoter sequences for investigating the evolutionary history of the studied promoters.

The second dataset, dataset_Ecoli, contains 11,884 artificially defined promoter sequences of Escherichia coli, along with the corresponding gene expression strengths measured by dRNA-seq. Since prokaryotes do not have many regulatory elements like eukaryotic enhancers, the expression level of their corresponding genes can be indirectly regarded as the strength of the promoter. This dataset originates from Thomason et al. [27] and was employed by Wang et al. [21] for predictive model. We also used dataset_Ecoli to train our model. The third dataset, dataset_trc, comprises 3,665 mutated trc promoter sequences and their corresponding promoter strengths. This dataset, introduced by Zhao et al. [29], was constructed using 83 rounds of mutation-constructionscreening-characterization engineering cycles. The strength of the promoters was determined by fluorescent protein intensity. We employed dataset_trc to validate the transfer learning capability of our constructed predictive model and test the predictive performance after fine-tuning.

### 4.2 Merged Chaos Game Representation

Merged Chaos Game Representation, abbreviated as Merged CGR, is a novel feature extraction method proposed for the first time in this paper. This approach is built upon the conventional Chaos Game Representation [24] and extends its applicability to not only the given sequence but also its related sequences, converting them into a unified matrix. Subsequently, this matrix is input into the DenseNet [25] for further processing. Notably, this representation method functions as an alternative to the widely employed position-specific scoring matrix (PSSM) found in other related studies.

For a sequence *s* = (*s*_1_*s*_2_ *. . . s_n_*), generating the corresponding Merged CGR involves three steps. The first step is to transform the sequence into a matrix using conventional CGR. The CGR sequence corresponding to *s*, *X_i_* = (*x_i_, y_i_*) where *i* = 1*, …, n*, is given by:

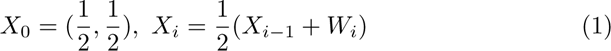

where *W_i_* equals to (0, 0), (1, 0), (0, 1), (1, 1) if *s_i_* is *A, T, C, G* respectively. By uniformly subdividing the unit square into *L*^2^ subsquares and calculating the number of points within each subdivision, the CGR sequence can be further transformed into a CGR matrix with *L* rows and *L* columns, denoted as *CGR*(*s*) (*L* = 20). The second step involves using BLAST [31] (blastn-short, evalue = 1) to search for matching sequences *s*^(1)^*, …, s*^(^*^m^*^)^ in dataset_pro. Subsequently, the Needleman-Wunsch algorithm [32] is applied to confirm their similarity to *s*. The similarity is calculated as the sum of the scores in the pairing, where both mismatch and gap have a score of -1, and match has a score of 1. This sum is then divided by the promoter length of 50, resulting in similarity scores *a*_1_*, …, a_m_* (*a_i_ ≤* 1). Finally, considering all sequences with similarity scores greater than 0, the Merged CGR matrix is computed as 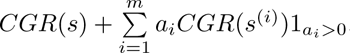.

We have chosen to apply Merged CGR instead of PSSM to extract the evolutionary information for promoters for the following reason. We aim to restrict our search for related sequences to the promoter region. Considering the relatively limited availability of promoter data, which frequently includes orphan promoters, it becomes challenging to identify highly similar sequences suitable for PSSM generation. In contrast, our Merged CGR method excels in integrating sequences exhibiting moderate or localized similarity, thereby accommodating such orphan promoter scenarios. Furthermore, a noteworthy advantage of our method lies in the resulting feature matrix, which takes the form of a square matrix as opposed to the elongated structure of the PSSM. This characteristic enables a more effective utilization of convolutional neural networks for feature extraction.

### 4.3 Word2vec Word Embedding

DNA sequences can be divided into a series of k-mers [52] [53] [54], allowing the sequence to be treated as text where the k-mers serve as words. Accordingly, we can use word embedding techniques from natural language processing (NLP) to represent these k-mers numerically.

The word2vec method, proposed by Mikolov et al. [55] [56], embeds words into meaningful high-dimensional numerical vectors. This is accomplished by conducting training on large textual data using shallow neural networks, thereby producing word embedding vectors within the hidden layer of the neural network. The fundamental principle behind word2vec is that words with similar contexts have similar semantics. The neural network structure includes the continuous bag-of-word (CBOW) model or the skip-gram model. During training, CBOW mainly predicts a word from its context, while skip-gram predicts the context words given a certain word. For instance, given a sequence of training words *w*_1_, *w*_2_, *w*_3_, . . . , *w_T_* , the objective of the skip-gram model is to maximize the average log probability described as:

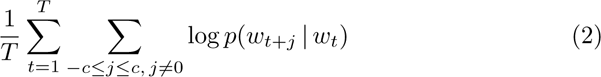

where *c* is the size of the training context (which can be a function of the center word *w_t_*). The probability is defined as:

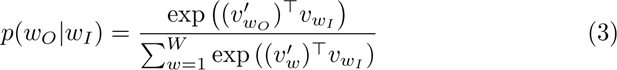

where *v_w_* and *v^′^* represent the ‘input’ and ‘output’ vector representations of *w* respectively, and *W* is the total number of words in the vocabulary. Each word’s vector representation during training is influenced by its surrounding vocabulary. If two words have similar contextual vocabulary, their word vectors will also be similar. Compared to the traditional one-hot embedding, word2vec offers denser representations and richer information for more profound semantic understanding and applications, making it more advantageous for training.

To better extract textual information from promoter sequences, we employed the word2vec method to obtain k-mer word embeddings in promoter sequences, following the specific strategies of the kmer2vec method [34]. Initially, we divided all promoter sequences in dataset_pro into a series of 3-mers using an overlapping division, treating them as complete text. Given a DNA sequence, for instance, AGGTAACAA (*n* = 9, length), using overlapping division, it can be decomposed into (*n k* + 1) k-mers (k=3), AGG GGT GTA TAA AAC ACA CAA. After a k-mer is taken, the starting position changes from *i* to *i* + 1 (*i* is the present position), and then the next k-mer will be taken.

Subsequently, we conducted training (window_size = 24; vector_size = 100; the skip-gram algorithm) on the text created from dataset_pro. This process provided word vectors corresponding to each 3-mer, reflecting both the sequence similarity of the 3-mers and their functional similarity in the evolutionary process of promoters. Then the significant contextual information in evolution was captured. During the training and testing phases of our model, for each promoter sequence, we fed the downstream modules with the numerical embeddings of its 3-mer vocabulary.

This method allows us to leverage the word2vec approach for representing k-mer sequences in DNA, capturing both sequence similarity and functional characteristics within the evolutionary context of promoters.

### 4.4 DenseNet

DenseNet (Densely Connected Convolutional Network) [25] is a deep learning algorithm designed for processing matrix data. It serves as an extension of traditional convolutional neural networks (CNN), with its most prominent feature being the dense connections between layers. These connections enable comprehensive information exchange among layers, allowing shallow-level information to be retained even after passing through multiple layers. This, in turn, allows for the training of deeper networks. Unlike ResNet [57], where connections involve addition, DenseNet employs a different approach by concatenating data across the channel dimension. Specifically, for the l-th layer, it takes the outputs of all preceding l-1 layers as inputs, that is,

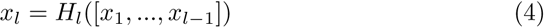

where *H_l_* is the function corresponding to the l-th layer. The channel output of *H_l_* assumes a crucial role in DenseNet, representing the growth rate of channels within the network (growth rate = 16 in this paper). Each layer in this study comprises four fundamental components. In addition to the standard components commonly found in CNNs, including the convolution part, Batch Normalization, and the activation part (ReLU), an additional dropout component [58] has been introduced to address overfitting.

### 4.5 Transformer

The Transformer [26] is a deep learning algorithm extensively employed in processing sequential data. Its core concept is the attention mechanism. Specifically, each element in sequential data, such as words, is transformed into vectors using techniques like word embedding. These vectors are further mapped to corresponding query (Q), key (K), and value (V) through various linear transformations. This allows each word to establish connections with all other words, as defined by the following equation:

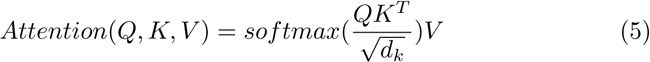

where *d_k_*is the dimension of Q and K. This attention mechanism is more capable of capturing longer-range information compared to traditional RNN-based algorithms [59][60]. Furthermore, the Transformer employs a multi-head strategy, linearly projecting the Q, K, V multiple times with different learned linear projections, enhancing the richness of information capture. In other words, we have

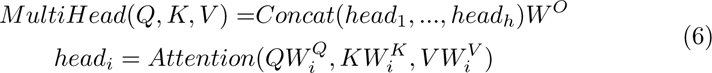

where *W^O^*is also a learned parameter matrix.

The core mechanism of the Transformer has been introduced above. In fact, the Transformer does not simply rely on this mechanism alone. It employs an encoder-decoder structure, with each part consisting of multiple Transformer blocks. Within each block, the Transformer combines the attention mechanism with residual networks, transforming the output into the input for the next block. In this paper, because we do not need to convert sequences into sequences, but only into real numbers, we have exclusively employed the encoder. The encoder consists of two Transformer blocks, with the number of attention heads set to 8.

### 4.6 Directed Evolution of Promoters In Silico

Using our model to perform promoter directed evolution requires the following two steps.

Step 1 (Random Mutation): First, we should choose the target promoter and its evolution direction. For example, in our project, we intended to evolve the promoter PnisA to have a stronger strength. If the sequence length is largar than 50 bp, we only consider the last 50 bp. We then used a program to randomly mutate the sequence of this promoter, and the number of mutated bases can be adjusted (usually <= 5). We can then obtain a large number of mutated sequences on the basis of the original promoter sequence.

Step 2 (Strength Screening): We input the large number of mutant sequences into our model and selected the top sequences with the highest predicted strength based on the model feedback. In this way, we completed the directed evolution of the promoter in silico, which can then be tested experimentally.

### 4.7 Bacterial Strains and Cultivation

The E. coli Stbl3 strain was used for plasmid cloning and fluorescent protein expression analysis. Inoculates were cultured in 5 mL of Luria Bertani (LB) medium containing antibiotics (kanamycin for the pET28A plasmid and carbenicillin for the pBV220 plasmid) in 15 mL shaking culture tubes at 200 rpm, 37℃. For gene expression analysis, with respect to constitutive promoters, we inoculated 50 µL of bacterial culture into 1 mL of LB medium containing antibiotics and incubated at 37°C with shaking at 200 rpm for 12-14 hours to ensure bacterial growth in the logarithmic phase. For the heat-inducible PL promoter, we initially inoculated 50 µL of bacterial culture into 1 mL of LB medium containing antibiotics and incubated at 37°C with shaking at 200 rpm for 4-6 hours. Subsequently, cells were induced expression by placing the culture in a 42°C water bath for 1 hour and then returned it to 37°C with shaking at 200 rpm for 10-12 hours. All strains and plasmids are listed in Supplementary Table 1.

### 4.8 Plasmid Construction

The PnisA promoter was ligated into the pET28a-GST-mCherry vector. Specific promoter mutation primers were designed for each promoter (Supplementary Table 2). PCR amplification was performed using Q5® High-Fidelity DNA Polymerase (NEB #M0491L). The original vector was digested with the restriction enzyme DpnI (NEB #R0176S) for 4 hours at 37°C. Subsequently, the amplified PCR fragments were separated by 1.5% agarose gel electrophoresis, and gel purification was carried out using the NucleoSpin·Gel and PCR cleanup kit (MN #740609.25). The PCR fragments and linearized pET28a-GST-mCherry vector were then subjected to homologous recombination using the NEBuilder® HiFi DNA Assembly Master Mix (NEB #E2621L) to obtain the ligated product. For the PL promoter, specific promoter mutation primers were designed (Supplementary Table 2). Q5® High-Fidelity DNA Polymerase (NEB #M0491L) was used for circular PCR on the pBV220-mCherry vector, resulting in the pBV220-PL-mCherry plasmid. The original vector was digested with the restriction enzyme DpnI (NEB #R0176S) for 4 hours at 37°C. The ligated products obtained were transformed into E. coli Stbl3 strain using the heat shock method. The sequence accuracy was confirmed through Sanger sequencing.

### 4.9 Fluorescence Expression Assay

For constitutive promoter PnisA, 50 µL of bacterial culture was inoculated at a 1:50 dilution into 2.5 mL LB medium containing kanamycin, and the culture was placed on a shaker at 200 rpm and 37°C for 14 hours to allow the cells to enter the logarithmic growth phase. For the temperature-inducible PL promoter, after reaching the logarithmic growth phase, the culture was induced using the method described above, followed by incubation on a shaker at 200 rpm and 37°C for an additional 10-12 hours. Subsequently, 0.5 mL of the bacterial culture was centrifuged at 4000 rpm for 5 minutes, the supernatant was discarded, and the pellet was resuspended in 1 mL of PBS buffer. After another centrifugation at 4000 rpm for 5 minutes and discarding the supernatant, the pellet was resuspended in 0.2 mL of PBS. The mCherry fluorescence and OD600 were then measured using a microplate reader.

### 4.10 Flow Cytometry

1. E. coli in the logarithmic growth phase were harvested, centrifuged at 4000 rpm for 5 minutes, and resuspended in PBS buffer. Flow cytometry data were collected using BD Fortessa or Thermo Fisher Attune NxT flow cytometers and analyzed with FlowJo software (BD Biosciences).

## 5 Declarations

### 5.1 Ethics Approval and Consent to Participate

Not applicable.

### 5.2 Availability of Data and Materials

The data presented in this study can be downloaded in the public database, and also available in Supplementary Materials. The source code implemented in this study can be found in our GitHub repository https://github.com/BobYHY/CAPE. Considering potential updates, please refer to our GitHub repository for instructions on accessing the website.

### 5.4 Funding

National Natural Science Foundation of China (NSFC) grant (12171275) Tsinghua University Education Foundation fund (042202008)

Tsinghua University Initiative Scientific Research Program

Academic and Scientific Works Competition for Undergraduates, Academic Affairs Office, Tsinghua University

Xuetang Program, Tsinghua University.

### 5.5 Authors’ Contributions

Conceptualization, R.R., H.Y., and S.Y.; Methodology, R.R., H.Y., and S.Y.; Investigation: R.R., J.T., and S.M.; Software: R.R., H.Y., Y.T., and Z.B.; Data curation, R.R., Z.B., H.Y. and Y.T.; Formal analysis: R.R., H.Y., and J.T.; Validation: R.R., J.T., S.M., H.Y. and Y.T.; Visualization: R.R., H.Y., and J.T.; Writing—original draft preparation, R.R., H.Y., and J.T.; Writing— review and editing, R.R. and S.Y.; Resources: S.Y. and R.R.; Supervision: S.Y.; Project Administration: S.Y.; Funding Acquisition: S.Y. and R.R. All authors have read and agreed to the published version of the manuscript.

## Supporting information

All Supplementary Materials

## Acknowledgments

Sincere thanks are extended to all members and advisors of Tsinghua iGEM 2022 for their dedicated efforts towards the incubation of this project. For the website of Tsinghua iGEM 2022, please refer to https://2022.igem.wiki/tsinghua/. Additionally, Professor Stephen S.-T. Yau is grateful to the National Center for Theoretical Sciences (NCTS) for providing an excellent research environment while part of this research was done. We are also grateful to Innovation Laboratory for Undergraduates, Center for Laboratory Training in Life Sciences (CLTLS), Tsinghua University.

