## Supplementary material for "Evolution is All You Need in Promoter Design and Optimization": All Supplementary Materials

### Supplementary Table1

| Plasmid | Source |
|---|---|
| pBV220-mCherry | ZOMANBIO |
| pET28a-GST-mCherry | ZOMANBIO |
| Materials | Catalogue |
| Stbl3 Chemically Competent Cell | AlpalifeBio #KTSM110L |
| NEBuilder® HiFi DNA Assembly Master Mix | NEB #E2621L |
| Q5® High-Fidelity DNA Polymerase | NEB #M0491L |
| NucleoSpin Gel and PCR clean-up | MN #740609.25 |
| DpnI | NEB #R0176S |
| Laboratory Apparatus | Manufacturer |
| Benchtop centrifuge | Eppendorf |
| PCR instrument | Eppendorf |
| Fortessa | BD |
| Microplate Reader | BIO-RAD 550 |

### Supplementary Table2

| Primer | |
|---|---|
| PnisA_F | tcgatcccgcgaaatctagtcttataactatactgacaatagaaacattaacaaatc |
| PnisA_pre_R | aagttaaacaaaattatttctagaggataatttattttgtagttccttcgaacgaaatcattgtatctaacaaact |
| PnisA01_R | aagttaaacaaaattatttctagaggataatttattttgtagtcccttctaactaaatcattgtatctaacaaact |
| PnisA02_R | aagttaaacaaaattatttctagaggataatttagtctgtagttccttcgtccgaaatcattgtatctaacaaact |
| PnisA03_R | aagttaaacaaaattatttctagaggataacttagtttgtagttccttctaactaaattattgtatctaacaaact |
| PnisA04_R | aagttaaacaaaattatttctagaggaaaatttattttgtagtcccttctaactaaatcattgtatctaacaaact |
| PnisA07_R | aagttaaacaaaattatttctagaggataacatagtttgtagttccttctaaccaaatcattgtatctaacaaact |
| PnisA10_R | aagttaaacaaaattatttctagaggatagttcattttgtagttcattctaacgaaatcattgtatctaacaaact |
| PnisA13_R | aagttaaacaaaattatttctagaggatagtttagtttgtagttccttctaactaaatcattgtatctaacaaact |
| PnisA14_R | aagttaaacaaaattatttctagaggataatctatcttgtagtcccttctaacaaaatcattgtatctaacaaact |
| PL01_F | cataaataccactggcggtcatacagagaacataagcagg |
| PL01_R | gttctctgtatgaccgccagtggtatttatgttaacaccgc |
| PL02_F | cataaatactactggcggtcatactgagaacaacagcagg |
| PL02_R | gttctcagtatgaccgccagtagtatttatgtcagcaccgc |
| PL03_F | cataactaccactggcggttatactgagtacatcagcagg |
| PL03_R | gtactcagtataaccgccagtggtagttatgtaaacaccgc |
| PL04_F | cataactaccactggcggtgatactaactacatcagcagg |
| PL04_R | gtagttagtatcaccgccagtggtagttatggcaacaccgc |
| PL05_F | cataaatactactggcggtgatactgagtacatcatcagg |
| PL05_R | gtactcagtatcaccgccagtagtatttatgtcaacatcgc |
| PL06_F | catcaataccactggcggtgagactgagtacatcatcaggacgcac |
| PL06_R | gtactcagtctcaccgccagtggtattgatgtcaacactgccagag |
| PL07_F | cataaataccactggcggttatactgagtacatcatcagg |
| PL07_R | gtactcagtataaccgccagtggtatttatgttaacgccgccagagata |
| PL08_F | cataaatactactagcggtaatactgagtacatcagcagg |
| PL08_R | gtactcagtattaccgctagtagtatttatgtaaacaccgc |
| PL09_F | cataaatactactggcggtgatactaagcacgttagcaggacg |
| PL09_R | gtgcttagtatcaccgccagtagtatttatgtcaacatcgc |
| PL10_F | cataaataccactggcggttatacagaggacataagcagaacgcac |
| PL10_R | gtcctctgtataaccgccagtggtatttatgtcaacaccgc |
| PL11_F | cataaataccactggcggtcctactgaacacataagcagg |
| PL11_R | gtgttcagtaggaccgccagtggtatttatgtccacaccgc |
| PL12_F | cataaataccactggcggttatactaagcacataaacaggacgcactg |
| PL12_R | gtgcttagtataaccgccagtggtatttatgtcaacatcgc |
| PL13_F | cgtaactaccactggcggttacactgagtacatcagcagg |
| PL13_R | gtactcagtgtaaccgccagtggtagttacgtcaacaccgc |
| PL14_F | catacatacaactggcggtgatacagagaacatcagcagg |
| PL14_R | gttctctgtatcaccgccagttgtatgtatgttaacaccgc |

#### Promoter Mutant Sequence

| PnisA | Predicted strength (log) |
|---|---|
| agtttgttagatacaatgattttgttagaatgaactacaaaatgaattgt | 8.632201195 |
| agttagttagatacaatgatgtcgttcgaaggaactacagactaaatgat | 8.632535934 |
| tgtttgttagatacaatgatttcgttagaagggactactaaataaaatat | 8.633365631 |
| agtatgttagatacaatgatttcgttccaagggactacaagataaatgat | 8.634530067 |
| agtttgttagatacaatgatttagttagaagggactacaaaataaattat | 8.636847496 |
| agtttgttagatacaatgatttagttagaagggactacaaaataaattat | 8.636847496 |
| agtttgttagatacaatgatttcggacgaaggaactacagactaaattat | 8.637728691 |
| agtttgttagatacaataatttagttagaaggaactacaaactaagttat | 8.645664215 |
| agtttgttagatacaatgatttagttagaagggactacaaaataaatttt | 8.655913353 |
| agtttgttagatacaatgatttagttagaagggactacaaaatagattat | 8.658157349 |
| agtttgttagatacaatgatttagttcgaaggaactacaagagaagttat | 8.658870697 |
| agtttgttagatacaatgatttggttagaaggaactacaaactatgttat | 8.659448624 |
| agttagttagatacaatgatttcgttagaagggactacaaaataaattgt | 8.671599388 |
| agtttgttagatacaatgatttagttagaagggactacaatataaattat | 8.685007095 |
| agtttgttagatacaatgatttcgttagaatgaactacaaaatgaactat | 8.687964439 |
| agttcgttagatacaatgagttcgttcgaaggaactgcaaactaacttat | 8.69769001 |
| agtttgttagatacaaagagttagttcgaaggaactacaaactagattat | 8.70376873 |
| agtttgttagatacaatgatttagttagaaggaactacaaactaaactat | 8.705487251 |
| tgtttgttagatacaatgatttagttagaagggactacaaaataaattat | 8.744571686 |
| agtttgttagatacaatgattttgttagaagggactacaagatagattat | 8.842524529 |
| previous | |
| agtttgttagatacaatgatttcgttcgaaggaactacaaaataaattat | 7.886957169 |

| PL | Predicted strength (log) |
|---|---|
| gcggtgttgacataaataccactggcggctatactaaccacataagcagg | 7.681978703 |
| gcggtgttgacataaataacactagcggttatactgagtacatcaggagg | 7.683638096 |
| gcggtgtgtacataaataccactggcggtgctattgagtacatcagcagg | 7.68473959 |
| gcggtgttgacataaatactactggcggtcatactaagcgcataagcagg | 7.68588829 |
| gcgatgttgacataactaccactggcggcgatactgagtacatcatcagg | 7.685947418 |
| gcggtgttaacataaataccactggcggtcatacagagaacataagcagg | 7.688251972 |
| gcggtgctgacataaatactactggcggtcatactgagaacaacagcagg | 7.689864635 |
| gcggtgtttacataactaccactggcggttatactgagtacatcagcagg | 7.690360546 |
| gcggtgttgccataactaccactggcggtgatactaactacatcagcagg | 7.694915771 |
| gcgatgttgacataaatactactggcggtgatactgagtacatcatcagg | 7.703474522 |
| gcagtgttgacatcaataccactggcggtgagactgagtacatcatcagg | 7.708014965 |
| gcggcgttaacataaataccactggcggttatactgagtacatcatcagg | 7.716670036 |
| gcggtgtttacataaatactactagcggtaatactgagtacatcagcagg | 7.716761589 |
| gcgatgttgacataaatactactggcggtgatactaagcacgttagcagg | 7.728304386 |
| gcggtgttgacataaataccactggcggttatacagaggacataagcaga | 7.739882946 |
| gcggtgtggacataaataccactggcggtcctactgaacacataagcagg | 7.750919819 |
| gcgatgttgacataaataccactggcggttatactaagcacataaacagg | 7.755888939 |
| gcggtgctaacataaataccactggcggttatactgagcacgtaagcagg | 7.75900507 |
| gcggtgttgacgtaactaccactggcggttacactgagtacatcagcagg | 7.767437458 |
| gcggtgttaacatacatacaactggcggtgatacagagaacatcagcagg | 7.775942802 |
| previous | |
| gcggtgttgacataaataccactggcggtgatactgagcacatcagcagg | 6.598791599 |

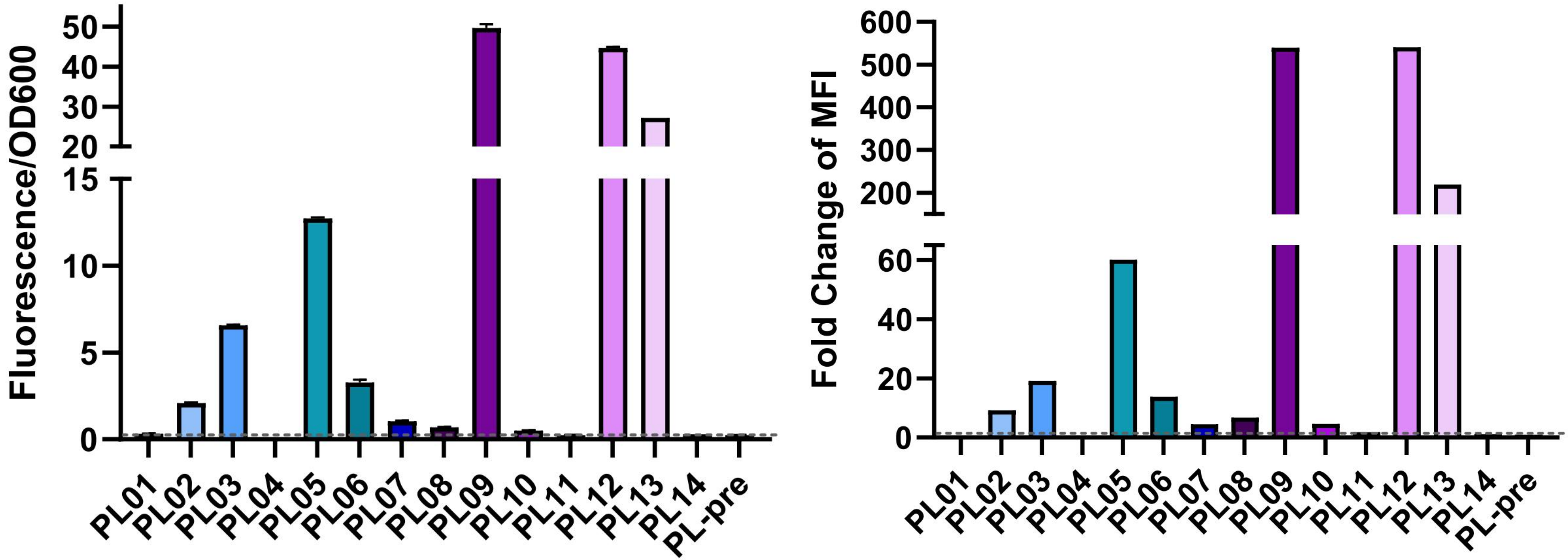

**Results of PL Before Inducing**
