## Supplementary figures and images for "Evolution is All You Need in Promoter Design and Optimization"

### Supplementary Figure 1

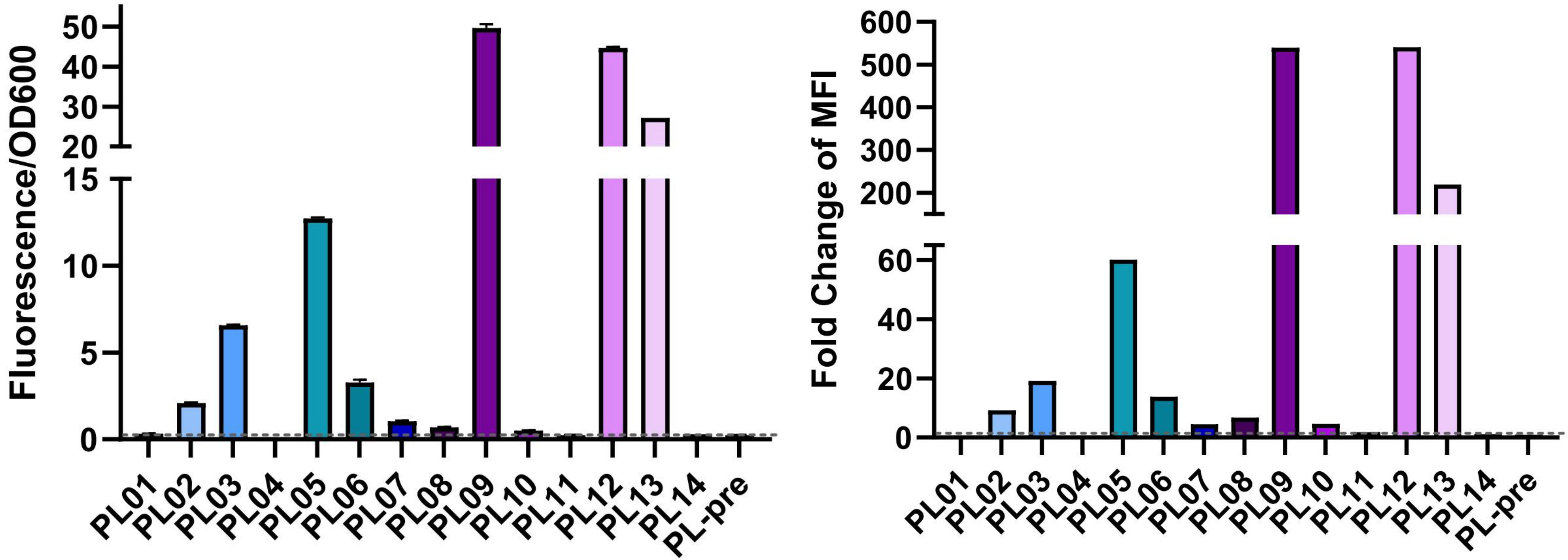


**Results of PL Before Inducing**
